# Identification of enriched hyperthermophilic microbial communities from a deep-sea hydrothermal vent chimney under electrolithoautotrophic culture conditions

**DOI:** 10.1101/2020.11.11.377697

**Authors:** G. Pillot, O. Amin Ali, S. Davidson, L. Shintu, A. Godfroy, Y. Combet-Blanc, P. Bonin, P.-P. Liebgott

## Abstract

Deep-sea hydrothermal vents are extreme and complex ecosystems based on a trophic chain. We are still unsure of the identities of the first colonizers of these environments and their metabolism, but they are thought to be (hyper)thermophilic autotrophs. Here we investigate whether the electric potential observed across hydrothermal chimneys could serve as an energy source for these first colonizers. Experiments were performed in a two-chamber microbial electrochemical system inoculated with deep-sea hydrothermal chimney samples, with a cathode as sole electron donor, CO_2_ as sole carbon source, and nitrate, sulfate, or oxygen as electron acceptors. After a few days of culture, all three experiments showed growth of electrotrophic biofilms consuming the electrons (directly or indirectly) and producing organic compounds including acetate, glycerol, and pyruvate. Within the biofilms, the only known autotroph species retrieved were members of *Archaeoglobales*. Various heterotrophic phyla also grew through trophic interactions, with *Thermococcales* growing in all three experiments as well as other bacterial groups specific to each electron acceptor. This electrotrophic metabolism as energy source driving initial microbial colonization of conductive hydrothermal chimneys is discussed.

## Introduction

Deep-sea hydrothermal vents are geochemical structures housing an extreme ecosystem rich in micro- and macro-organisms. Since their discovery in 1977 (Corliss and Ballard, 1977), they have attracted the interest of researchers and, more recently, industries with their unique characteristics. Isolated in the deep ocean, far away from sunlight and subsequent organic substrates, the primary energy sources for the development of this abundant biosphere remain elusive in these extreme, mineral-rich environments. Since their discovery, many new metabolisms have been identified based on organic or inorganic molecules. However, the driving force sustaining all biodiversity in these environments is thought to be based on chemolithoautotrophy (Alain *et al.*, 2004). Unlike most ecosystems, deep-sea ecosystems are totally dark, and microorganisms have thus adapted to base their metabolism on lithoautotrophy using inorganic compounds as the energy source to fix inorganic carbon sources.

Primary colonizers of deep-sea hydrothermal vents are assumed to be (hyper)thermophilic microbes developing near hydrothermal fluid, as retrieved in young hydrothermal chimneys. These first colonizers are affiliated to *Archaea*, such as *Archaeoglobales*, *Thermococcales, Methanococcales* or *Desulfurococcales*, and to *Bacteria*, for example ε-*proteobacteria* and *Aquificales* (Huber *et al.*, 2002; Nercessian *et al.*, 2003; Takai *et al.*, 2004). Recent studies have also shown that hyperthermophilic *Archaea*, which count among the supposed first colonizers, are able to quickly scan and fix onto surfaces to find the best conditions for growth (Wirth *et al.*, 2018). These hyperthermophilic microorganisms fix inorganic carbon through chemolithoautotrophic types of metabolism, using H_2_, H_2_S or CH4 as energy sources and oxidized molecules such as oxygen, sulfur compounds, iron oxide or even nitrate as electron acceptors.

However, the discovery of the presence of an abiotic electric current across the chimney walls (Yamamoto *et al.*, 2018) prompted the hypothesis of a new type of microorganism called “eletrotrophs”, microorganisms which have the capacity to use electrons from the abiotic electric current as an energy source coupled with carbon fixation from CO_2_. This metabolism was identified a few years ago on a mesophilic chemolithoautotrophic Fe(II)-oxidizing bacterium, *Acidithiobacillus ferrooxidans* (Ishii *et al.*, 2015). The mechanism of energy uptake from electrodes has been discussed since the discovery of biofilms growing on cathodes, and little is known about this process, unlike that of anodic electron transfer. The two main hypotheses are the use of similar direct electron transfer pathway as on the anode (Pous *et al.*, 2016), or the use of free cell-derived enzymes, which can interact with electrode surfaces to catalyze the electron transfers (Deutzmann *et al.*, 2015). Recent studies have shown the exoelectrogenic ability of some hyperthermophilic microorganisms isolated from deep-sea hydrothermal vents belonging to *Archaeoglobales* and *Thermococcales* (Pillot, *et al.*, 2018; Yilmazel *et al.*, 2018; Pillot *et al.*, 2019), but no studies have been done on environmental samples potentially harboring electrotrophic communities growing naturally with an electric current as their sole energy source.

In this article, we investigate the potential presence of electrotrophic communities in deep-sea hydrothermal vents capable of using electrons directly or indirectly from the abiotic current. To this end, we mimiced the conductive surface of the hydrothermal chimney in a cathodic chamber of Microbial Electrochemical Systems (MESs) with a polarized cathode to enrich the potential electrotrophic communities inhabiting these extreme environments. The polarized cathode served as the sole energy source, while CO_2_ bubbling served as sole carbon source. Under these experimental conditions, the distinct effects of the presence of nitrate, oxygen, and sulfate on the community taxonomic composition were investigated.

## Results

### Current consumption from electrotroph enrichments

Hydrothermal vents chimney samples were inoculated in MESs filled with sterile mineral medium and incubated at 80°C to enrich electrotrophic communities. The electrode (cathode) and the sparged CO_2_ were used as sole energy donor and carbon source, respectively. Nitrate, sulfate and oxygen were separately tested as electron acceptor. The electrode potential has been chosen in order to prevent the abiotic cathodic reduction of the culture medium (Fig. S1), thus the electrode was poised at −590 mV/SHE in the presence of sulfate and nitrate while this potential was −300 mV/SHE in the presence of oxygen. A fourth experiment with sulfate as electron acceptor at −300 mV/SHE provides information about the effect of the cathode potential on the microbial community. For comparison, microbial growth was also monitored in an inoculated system without any poised electrode during a month in the same incubation conditions. Interestingly, in the latter condition, without an electrode as energy source, no microbial growth occurred, a finding supported by microscope and spectrophotometric observations (data not shown). Moreover, no organic compounds were produced, supported by the HPLC and NMR measurements.

When the electrode was poised, abiotic controls containing no inoculum displayed constant currents of ≈0,016 A.m^−2^ at −590 mV and ≈0,01 A.m^−2^ at −300 mV/SHE. In both conditions, no hydrogen production on the cathode via water electrolysis, continuously monitored by μGC, was detected. The detection threshold of the μGC (>0.001% of total gas) indicated a theoretical production lower than 34 μM day-1 (data not shown) as previously reported at 25°C (Marshall *et al.*, 2012, 2013). In comparison, experiments with the chimney sample showed current consumptions increasing shortly after inoculation (Fig. 1). Indeed, when subtracting abiotic current, the current consumptions reached a stabilized maximum of 0.36 A.m^−2^ on oxygen, 0.72 A.m^−2^ on nitrate, and up to 1.83 A.m^−2^ on sulfate at −590mV or 1.38 A. m^−2^ on sulfate at −300 mV/SHE (data not shown), corresponding to 36, 45, 114 and 83-fold increases compared to abiotic current, respectively. MESs were then autoclaved, displaying decreased currents that were similar to the values of abiotic controls with a stabilized current around ≈0,021 A.m^−2^, indicating the importance of the living biofilm in the current consumption.

**Figure 1:**
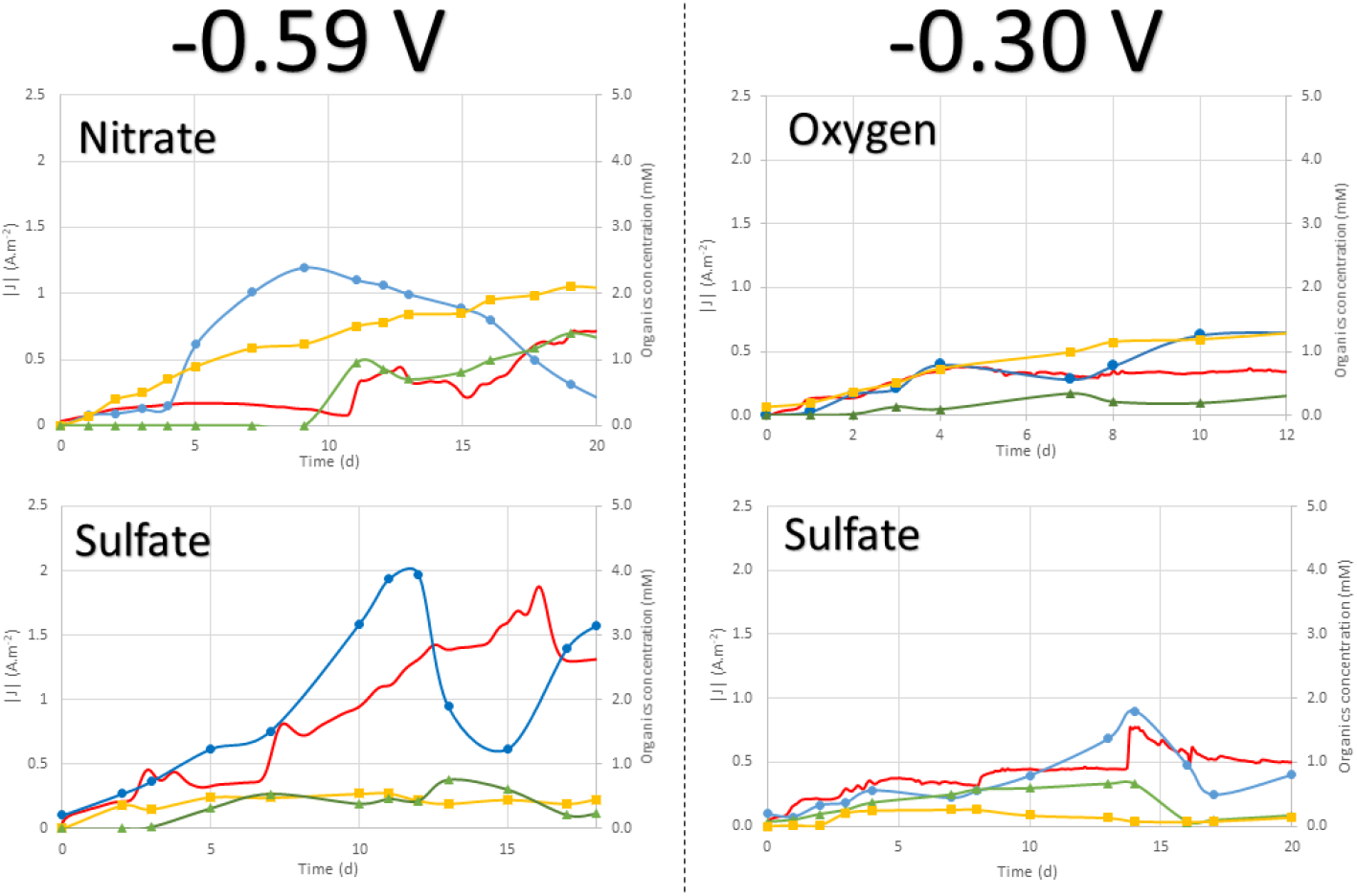
Current consumption (red continuous line); pyruvate (blue triangle), glycerol (yellow square) and acetate (green cross) productions over time of culture for each electron-acceptor experiment. The current was obtained from a poised electrode at −590 mV vs SHE for nitrate and sulfate experiments and −300 mV vs SHE for oxygen.

At the end of monitoring of current consumption, CycloVoltamograms (CVs) were performed to study reactions of oxidation and reduction that could occur in MESs (Fig. S2A). A first peak of reduction is observed at −0.295, −0.293 and −0.217 V vs SHE in presence of nitrate, sulfate and oxygen as electron acceptor, respectively (Fig. S2B). A second peak is observed at −0.639 and −0.502 V vs SHE on nitrate and sulfate, respectively. The on-set potential of H_2_ evolution was measured in our conditions at −0.830, −0.830, −0.780 and −0.830 V vs SHE in presence of nitrate, sulfate and oxygen as electron acceptor and in abiotic condition, respectively. No redox peaks were detected in the abiotic controls and freshly inoculated MESs, hence indicating a lack of electron shuttles brought with the inoculum (Fig. S2A).

### Organic compounds production in liquid media

During enrichment of the electrotrophic communities, the production of organic compounds was monitored in the liquid media. Data in presence of Nitrate, Oxygen and Sulfate at −300 mv and/or −590mV vs SHE are presented in Fig. 1. Interestingly, glycerol, pyruvate, and acetate were the dominant products released in all experimental runs. Glycerol increased slowly throughout the experiments to reach a maximum of 0.47 mM on sulfate (Day 11), 1.32 mM on oxygen (Day 12) and 2.32 mM on nitrate (Day 19). Acetate accumulated in the medium to reach 0.33 mM on oxygen (Day 7), 0.75 mM on sulfate (Day 13) and 1.40 mM on nitrate (Day 19). Pyruvate was exponentially produced after a few days of culture, reaching a maximum of 1.32, 2.39, and 3.94 mM in presence of oxygen (Day 12), nitrate (Day 9), and sulfate (Day 11), respectively. Pyruvate levels varied thereafter, due probably to microbial consumption or thermal degradation. Coulombic efficiency calculated on the last day of the experiment (Fig. 2) showed up to 71% (on nitrate), 89% (on oxygen) and 90% (on sulfate) of electrons consumed were converted to organic compounds and released into the liquid media. The rest represents the share of electrons retained in non-accumulated compounds (Table S1) and in the organic matter constituting the cells of the electrotrophic communities (estimated by qPCR to total between 10^8^ and 10^10^ 16S rRNA gene copies per MES Fig. 3).

**Figure 2:**
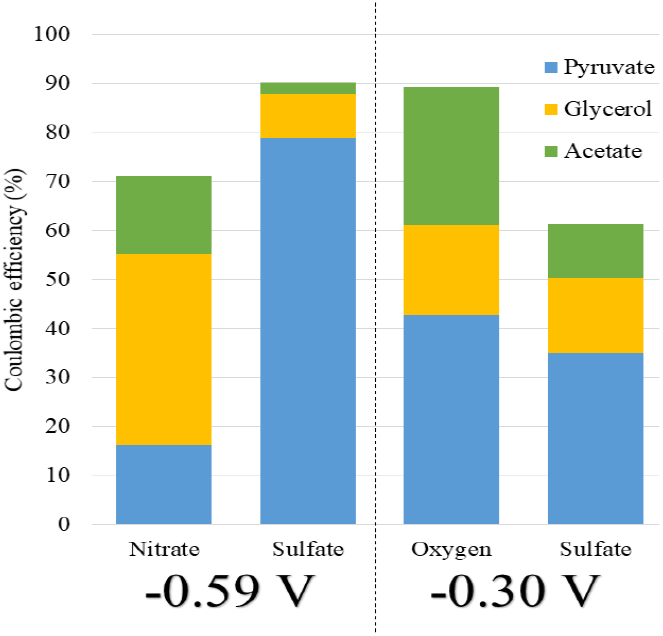
Coulombic efficiency for organic products in presence of the different electron acceptors and potentials.

**Figure 3:**
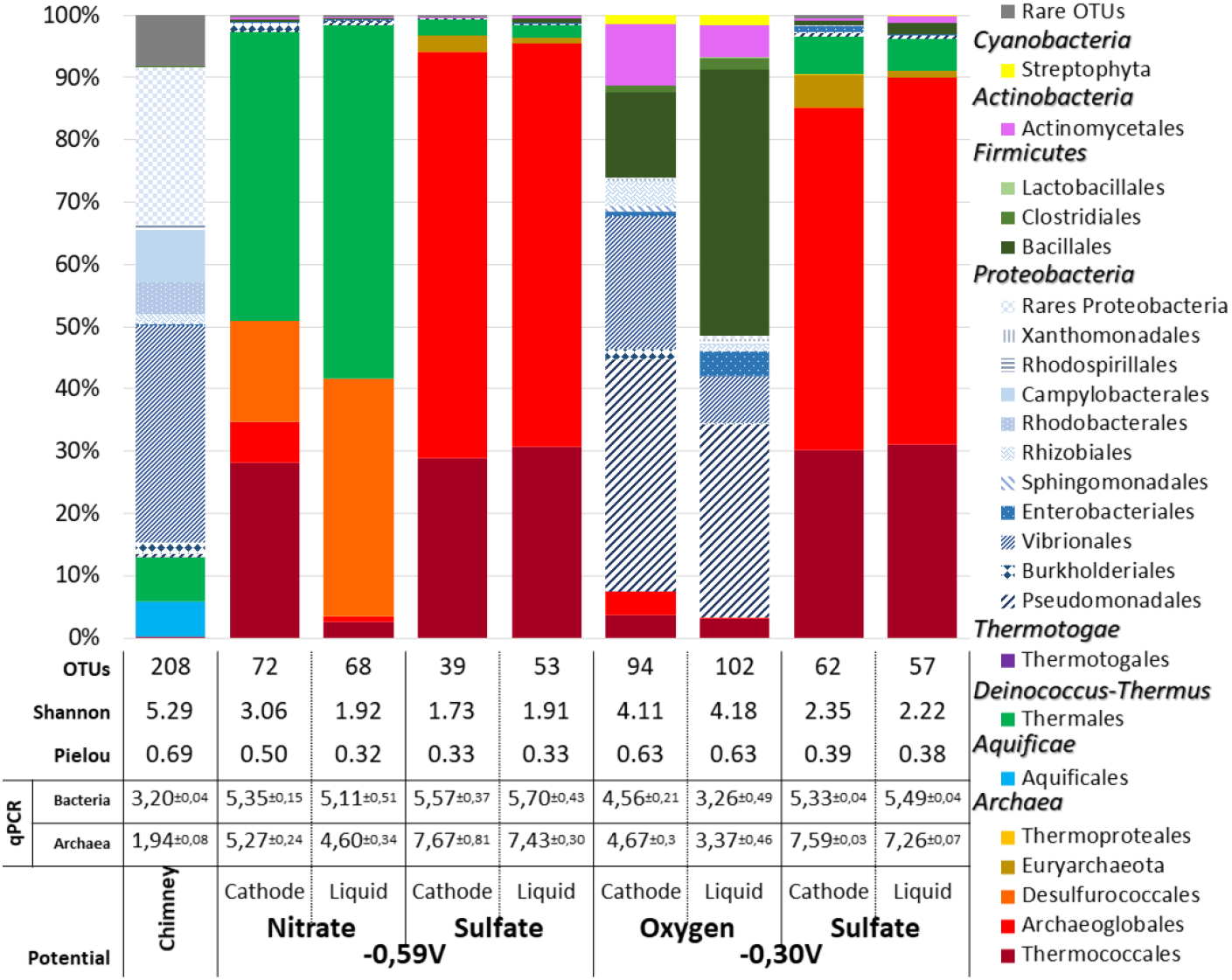
Dominant taxonomic affiliation at order level, biodiversity indices and qPCR of microbial communities from a crushed chimney sample from Capelinhos vent site (Lucky Strike hydrothermal vent field), as plotted on the cathode and liquid media (LM) after the weeks of culture. OTUs representing less than 1% of total sequences of the samples are pooled as ‘Rare OTUs’. qPCR expressed in Log10 Cells per gram of crushed chimney, per milliliter of liquid or per cm^2^ of working electrode. Standard deviation obtained on analytical triplicates.

### Biodiversity of electrotrophic communities on different electron acceptors

Once current consumption reached a stabilized maximum, DNA from the biofilm and from planktonic cells in culture media was extracted and sequenced on the V4 region of the 16S rRNA to study relative biodiversity. Figure 3 reports the taxonomic affiliation of the OTUs obtained. The chimney fragment inoculum showed a rich biodiversity (Shannon index at 5.29 and Pielou’s index at 0.69), with 208 OTUs mainly affiliated to *Bacteria* (99.49% vs 0.51% of *Archaea)* and more particularly to *Proteobacteria* from *Vibrionales* (34.8%), miscellaneous rare *Proteobacteria* (>33%), *Campylobacterales* (8.3%), *Thermales* (7.1%), *Aquificales* (5.62%), and *Rhodobacterales* (5.1%).

Enrichments in MES showed less biodiversity on cathodes and in liquid media, suggesting the selective development of functional communities. The Shannon index values were 3.1 and 1.9 on nitrate, 1.7 and 1.9 on sulfate, and 4.1 and 4.2 on oxygen, with fewer OTUs associated to 72 and 68 OTUs on nitrate, 39 and 53 on sulfate, and 94 and 102 on oxygen, all on electrodes and in liquid media, respectively. The taxonomic composition of these communities showed a larger proportion of *Archaea*, with 51% and 41.6% on nitrate, 96.8% and 96.3% on sulfate, and 7.6% and 3.4% on oxygen on the electrodes and in liquid media, respectively. Whatever the electron acceptor used, the archaeal population on the electrode was mainly composed of *Archaeoglobales* and *Thermococcales* at different relative abundances. These were present at proportions ranging from 6.7% and 28.2% on nitrate to 65.8% and 28.6% on sulfate and 3.8% and 3.6% on oxygen, respectively. Equivalent proportions of *Archaeoglobales* and *Thermococcales* were retrieved in liquid media, at 1.0% and 2.5% on nitrate, 65.8% and 29.6% on sulfate and 0.3% and 2.7% on oxygen, respectively (see Fig. 3). The MiSeq Illumina results served to study only 290 bp of 16S rRNA and thus to affiliate microorganisms confirmed at the family level, but they can also provide some information on the enriched genera.

To obtain more information on the probable *Archaeoglobales* and *Thermococcales* genus, we attempted a species-level identification through phylogenetic analysis. The results are presented in Fig. 4 as a Maximum likelihood phylogenetic tree. The dominant OTUs on sulfate and oxygen were closest to *Ferroglobus placidus* and *Archeoglobus fulgidus* (97.61%), whereas the dominant OTU on nitrate was affiliated to *Geoglobus ahangari* with an identity of 98.63%. The remaining part of the biodiversity was specific to each electron acceptor used. Enrichment on nitrate showed 13.8% and 28% of *Desulfurococcales* and 46.2% and 56.5% of *Thermales* on the electrode and in liquid media, respectively. Among *Thermales* that developed on the electrode, 30% were represented by a new taxon (OTU 14 in Fig. 4 and S3) whose closest cultured species was *Vulcanithermus mediatlanticus* (90% similarity). On sulfate, the remaining biodiversity represented less than 4% of the population but was mainly represented by two particular OTUs. The first OTU was affiliated to a new *Euryarchaeota* and accounted for up to 2.4% and 0.8% of the total population on the electrode and in liquid media, respectively (OTU 10 in Fig. 4 and S3). The closest cultured match (86% similarity) was *Methanothermus fervidus* strain DSM 2088. The second OTU was affiliated to the new *Deinococcales* species (< 97% similarity) and accounted in nitrate enrichment for 2.0% and 1.9% of the biodiversity on the electrode and in liquid media, respectively (OTU 14 in Fig. 4 and S3). In the enrichment on oxygen, the communities were dominated by 36.6% and 30.2% of *Pseudomonadales* (Pseudomonas sp.), 14% and 42.6% of *Bacillales (Bacillus* and *Geobacillus* sp.), 21.3% and 7.41% of *Vibrionales* (*Photobacterium* sp.), and 9.8% and 5.1% of *Actinomycetales* (spread across 9 species) on the electrode and in liquid media, respectively. The remaining biodiversity was spread across *Proteobacteria* orders.

**Figure 4.**
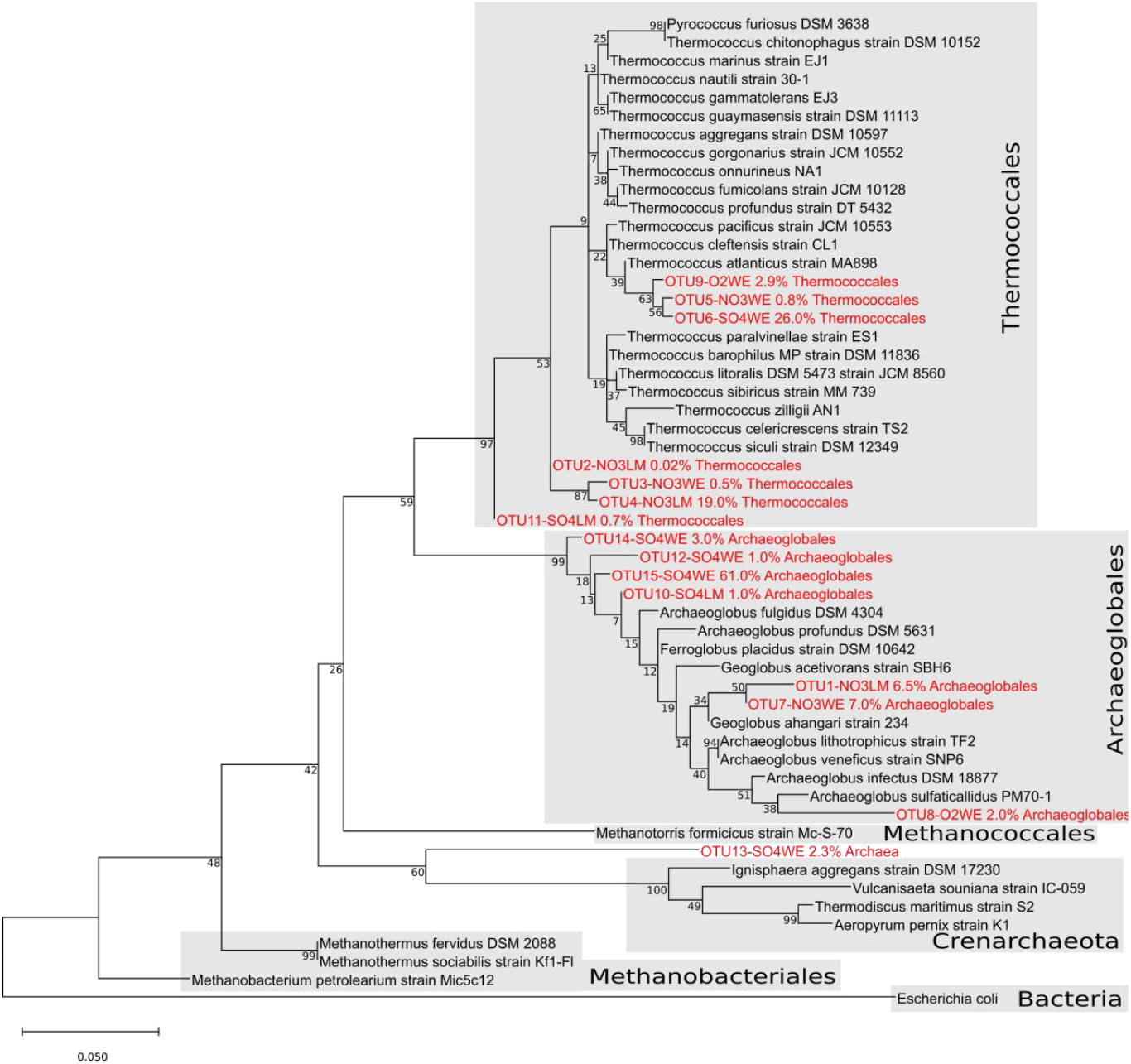
Maximum Likelihood phylogenetic tree of archaeal OTUs retrieved on various enrichments on the 293pb 16S fragment obtained in the barcoding 16S method (LM: Liquid Media; WE: Working Electrode, cathode). Numbers at nodes represent bootstrap values inferred by MEGAX. Scale bars represent the average number of substitutions per site.

The clustering of the dominant OTUs (at a threshold of 0.05% of total sequences) obtained previously on the chimney sample and enrichments in MES showed a clear differentiation of communities retrieved in each sample (Fig. S3). The Pearson method used on OTU distribution produced four clusters, one corresponding to the inoculum and the three others to each electron acceptor. Indeed, only two OTUs (OTU 4 and 36) were clearly shared between two different communities, one affiliated to *Thermococcus* spp. on nitrate and sulfate and one to *Ralstonia* sp. on the chimney sample and on nitrate. It is surprising to observe a recurrence of this last OTU, which could be a contaminant specific to the extraction kit used (Salter *et al.*, 2014). The other 50 dominant OTUs were specific to one community, with 21 OTUs on oxygen, 4 on sulfate, 8 on nitrate, and 17 on the chimney sample. The electrotrophic communities colonizing the cathode were therefore different depending on electron acceptor used and their concentration was too low to be detected in the chimney sample.

## Discussion

### Archaeoglobales as systematic (electro)litho-autotrophs of the community

We have evidenced the development of microbial electrotrophic communities and metabolic activity supported by current consumption (Fig.1), product production (Fig. 2), and qPCRs (Fig. 3). These data suggest that growth did occur from energy supplied by the cathode. Our study is the first to show the possibility of growth of biofilm from environments harboring natural electric current in the total absence of soluble electron donors. To further discuss the putative mechanism, it is necessary to have a look at our conditions unfavorable for water electrolysis (see Fig. S2). The equilibrium potential for water reduction into hydrogen at 80°C, pH 7, and 1 atm was calculated at −0.490 V vs SHE in pure water. The operational reduction potential is expected to be lower than the theoretical value due to internal resistances (from electrical connections, electrolytes, ionic membranes, etc.) (Lim *et al.*, 2017) and overpotentials (electrode material). This was confirmed with the on-set potential of H_2_ evolution measured at −0.830mV vs SHE in both experimental condition and abiotic control, indicating the absence of catalytic effect of putative hydrogenases secreted by the biofilm or metals from inoculum. Also, during preliminar potentials screening, the increase in current consumption and H_2_ production was observed only below −0.7V vs SHE (Figs. S1 and S2). Moreover, we obtained similar biodiversity on sulfate and oxygen with the cathode poised at −300 mV vs SHE (Fig. 3), 190mV more positive than the Equilibrium potential of H_2_ evolution, with then no electrochemical possibility of H_2_ production, even at molecular level. In addition, the presence of catalytic waves observed by CV with midpoint potentials between −0.217 V to −0.639 V indicate the implication of enzymes directly connected to the surface of the electrode (see Fig. S2). Finally, the fixation of 267 to 1596 Coulombs.day^−1^ into organics (Fig. 1) exceeds the maximum theoretical abiotic generation of hydrogen (~3 C.day^−1^) 90- to 530-fold.

Therefore, under our experimental conditions, the biofilm growth should be largely ensured by a significant part of a direct transfer of electrons from the cathode, thereby demonstrating the presence of electrolithoautotroph microorganisms.

Taxonomic analysis of the enriched microbial communities at the end of the experiments showed the systematic presence of *Archaeoglobales* on cathodes. Moreover, the qPCR and MiSeq data (Fig. 3) highlighted a strong correlation between current consumption and density of *Archaeoglobales* on the electrode (Fig. S4, R^2^=0.945).

The OTUs were related to some *Archaeoglobales* strains with 95-98% identities. Thus, we assume that under our experimental conditions new specific electrotrophic metabolisms or new electrolithoautotrophic *Archaeoglobaceae* species were enriched on the cathode. They were retrieved in all conditions and belonged to the only order in our communities exhibiting autotrophic metabolism. Autotrophic growth in the *Archaeoglobales* order is ensured mainly through using H_2_ as energy source and requires both branches of the reductive acetyl-CoA/Wood-Ljungdahl pathway for CO_2_ fixation (Vorholt *et al.*, 1997). Terminal electron acceptors used by this order include sulfate, nitrate, poorly crystalline Fe (III) oxide, and sulfur oxyanions (Brileya and Reysenbach, 2014). Moreover, *Archaeoglobus fulgidus* has been recently shown to grow on iron by directly snatching electrons under carbon starvation during the corrosion process (Amin Ali *et al.*, 2020). Furthermore, *Ferroglobus* and *Geoglobus* species were shown to be exoelectrogens in pure culture in a microbial electrosynthesis cell (Yilmazel *et al.*, 2018) and have been enriched within a microbial electrolysis cell (Pillot, *et al.*, 2018; Pillot *et al.*, 2019). Given these elements, the identified *Archaeoglobales* species could be, under our electrolithoautotrophic conditions, the first colonizers of the electrode during the first days of growth.

The growth of *Archaeoglobales* species in presence of oxygen is a surprising finding. *Archaeoglobales* have a strictly anaerobic metabolism, and the reductive acetyl-CoA pathway is very sensitive to the presence of oxygen (Fuchs, 2011). This can be firstly explained by the low solubility of oxygen at 80°C. Secondly, carbon cloth mesh reduces oxygen in the environment, allowing for anaerobic development of microorganisms into a protective biofilm (Hamilton, 1987). This observation was supported by the near absence of *Archaeoglobales* in the liquid medium (Fig. 3). One of the hypotheses concerns direct interspecies electron transfer (DIET) (Kato *et al.*, 2012; Lovley, 2017), with *Archaeoglobales* transferring electrons to another microorganism as an electron acceptor. Research into DIET is in its early stages, and further investigations are required to better understand the diversity of microorganisms and the mechanism of carbon and electron flows in anaerobic environments (Lovley, 2017) such as hydrothermal ecosystems.

### Electrosynthesis of organic compounds

Accumulation of pyruvate, glycerol and acetate was measured, while another set of compounds that appeared transiently were essentially detectable in the first few days of biofilm growth (Table S1). They included amino acids (threonine, alanine) and volatile fatty acids (formate, succinate, lactate, acetoacetate, 3-hydroxyisovalerate) whose concentrations did not exceed 0.1 mM. Despite their thermostability, this transient production suggests they were used by microbial communities developing on the electrode in interaction with the primary producers during enrichment.

On the other hand, in presence of nitrate, sulfate and oxygen as electron acceptors, the liquid media accumulated mainly acetate, glycerol, and pyruvate (Fig. 1). Coulombic efficiency calculations (Fig. 2) showed that electron content of the carbon products represented 71%–90% of electrons consumed, the rest being potentially used directly for biomass or transferred to an electron acceptor. This concurs with the energy yield from the Wood-Ljungdahl pathway of *Archaeoglobales*, with only 5% of carbon flux directed to the production of biomass and the other 95% diverted to the production of small organic end-products excreted from the cell (Fast and Papoutsakis, 2012).

Pyruvate is a central intermediate of CO_2_ uptake by the reducing pathway of the acetyl-CoA/WL pathway (Berg *et al.*, 2010). It can be used to drive the anabolic reactions needed for biosynthesis of cellular constituents. Theoretically, the only explanation for improved production and accumulation of pyruvate (up to 5 mM in the liquid media of sulfate experiment) would be that pyruvate-consuming enzymes were inhibited or that pyruvate influx exceeded its conversion rate. Here we could suggest that in-cell electron over-feeding at the cathode leads to significant production of pyruvate when the electron acceptor runs out.

In an ecophysiological context, similar pyruvate and glycerol production could occur on hydrothermal chimney walls into which electric current propagates (Yamamoto *et al.*, 2017). The electrotroph biofilms would continually receive electrons, leading to an excess of intracellular reducing power which would be counterbalanced by overproduction of glycerol and pyruvate (Björkqvist *et al.*, 1997; Furdui and Ragsdale, 2000). Furthermore, these products can serve as carbon and energy sources for heterotrophic microorganisms or for fermentation. In our experiments, pyruvate and glycerol concentrations varied over time, suggesting they were being consumed by heterotrophic microorganisms. Acetate production would thus result from the fermentation of pyruvate or other compounds produced by electrotrophic *Archaeoglobales*.

### Enrichment of rich heterotrophic biodiversity from electrotrophic Archaeoglobales community

During our enrichment experiments, the development of effective and specific biodiversity was dependent on the electron acceptors used (Fig. 3). Heatmap analyses (Fig. S3) showed four distinct communities for the three electron acceptors and the initial inoculum. Thus, at the lower taxonomic level of the biodiversity analysis, most OTUs are not common to multiple enrichments, except for one OTU of *Thermococcales* that was found in both the nitrate and sulfate experiments. This suggests a real specificity of the communities and a specific evolution or adaptation of the members of the shared phyla to the different electron acceptors available in the environment. However, the various enrichments also showed the presence of *Thermococcales* regardless of the electron acceptors used, thus demonstrating a strong interaction between *Thermococcales*, assumed to be heterotrophs, and *Archaeoglobales*, the only demonstrated autotrophs. Moreover, members of these two groups have frequently been found together in various hydrothermal sites (Corre et al., 2001; Nercessian et al., 2003; Takai et al., 2004; Jaeschke et al., 2012), where they are considered potential primary colonizers (Reysenbach *et al.*, 2000; Schrenk *et al.*, 2003, 2013; Lin *et al.*, 2016; Wirth, 2017). After *Thermococcales*, the rest of the heterotrophic biodiversity was specific to each electron acceptor.

On nitrate, two additional phylogenetic groups were retrieved: *Desulfurococcales* and *Thermales*. OTUs of *Desulfurococcales* are mainly affiliated to *Thermodiscus* or *Aeropyrum* species, which are hyperthermophilic and heterotrophic *Crenarchaeota* growing by fermentation of complex organic compounds or sulfur/oxygen reduction (Huber and Stetter, 2015). Concerning *Thermales*, a new taxon was enriched on cathode and only affiliated to *Vulcanithermus mediatlanticus* with similarity of 90 %. This new taxon of *Thermales* (OTU 14, Fig. S3) was also enriched up to 2% on the cathode of sulfate enrichment. *Thermales* are thermophilic (30°C–80°C) and heterotrophic bacteria whose only four genera (*Marinithermus*, *Oceanithermus*, *Rhabdothermus*, and *Vulcanithermus*) are all retrieved in marine hydrothermal systems. They can grow under aerobic, microaerophilic and some anaerobic conditions with several inorganic electron acceptors such as nitrate, nitrite, Fe (III) and elemental sulfur (Albuquerque and Costa, 2014). All of the *Thermales* species can utilize the pyruvate as carbon and energy source with the sulfate or nitrate as electron acceptors.

*Pseudomonadales* and *Bacillales* were found in the oxygen experiment. Most *Pseudomonas* are known to be aerobic and mesophilic bacteria, with a few thermophilic species (up to 65°C) (Lyons *et al.*, 1984; Palleroni, 2015). There have already been some reports of mesophilic *Pseudomonas* species growing in thermophilic conditions in composting environments (Droffner *et al.*, 1995). Moreover, some *Pseudomonas* sp. are known to be electroactive in microbial fuel cells through long-distance extracellular electron transport (Shen *et al.*, 2014; Maruthupandy *et al.*, 2015; Lai *et al.*, 2016), and were dominant on the cathodes of a benthic microbial fuel cell on a deep-ocean cold seep (Reimers *et al.*, 2006). In *Bacillales*, the *Geobacillus* spp. and some *Bacillus* sp. are known to be mainly (hyper)thermophilic aerobic and heterotrophic *Firmicutes* (Vos, 2015).

### Hydrothermal electric current: a new energy source for the development of primary producers

The presence of so many heterotrophs in an initially autotrophic condition points to the hypothesis of a trophic relationship inside the electrotrophic community (Fig. 5). This suggests that the only autotrophs retrieved in all communities, the *Archaeoglobales*, might be the first colonizer of the electrode, using CO_2_ as carbon source and the cathode as energy source. Models based on electron donor acceptor availability predicted low abundances of *Archaeoglobales* (<0.04%) due to low concentration of H_2_ (Lin *et al.*, 2016) whereas *in-situ* detection found abundances of more than 40% in the inner section of the studied hydrothermal chimney (Dahle *et al.*, 2018). The authors concluded on a probable H_2_ syntrophy, with hydrogen being produced by heterotrophic microorganisms such as fermentative *Thermococcales* species. Our study showed that Archaeoglobales can also grow electrolithoautotrophically, feeding on the natural electric current through the chimney walls. This new energy source, which is not considered in the models, would gap this model/observation difference and raises the question of the importance of this metabolism in the primary colonization of hydrothermal vents. It would allows a long-range transfer between the electron donor (H_2_S oxidized on the inner surface of the chimney wall) and the electron acceptors (O_2_, sulfur compounds, nitrate, metals) covering the external surface of the chimney. This electrical current would thus allow primary colonizers to grow, releasing organic compounds which are then used by the heterotrophic community, as observed in our experiments. Moreover,this would provide a constant source of electron donor to all over the surface of the chimney, allowing to meet a wider range of physiological conditions through pH, temperature, and oxidoreduction gradients. This allows a wider diversity of growth patterns than through chemolithoautotrophy, which is restricted to unstable and limited contact zones between reduced compounds (H_2_, H_2_S) in the hydrothermal fluid and electron acceptors around the hydrothermal chimneys (O_2_, SO_4_, NO_3_), which often precipitate together.

## Conclusion

Taken together, the results found in this study converge into evidence of the ability of indigenous microorganisms from deep hydrothermal vents to grow autotrophically (CO_2_ used as only carbon source) using electric current as energy source. This ability seems to be spread across diverse phylogenetic groups and to be coupled with diverse electron acceptors. Through their electro-litho-auto-trophic metabolism, *Archaeoglobaceae* strains could produce and release organic compounds into their close environment, allowing the growth of heterotrophic microorganisms and ultimately enabling more and more diversity to develop over time (Fig. 5). This metabolism could be one of the primary energies for the colonization of deep-sea hydrothermal chimneys and the development of a complex trophic network driving sustainable biodiversity. A similar mechanism could have occurred during the Hadean, allowing the emergence of life in hydrothermal environments through constant electron influx to the first proto-cells.

## Experimental procedures

### Sample collection and preparation

A hydrothermal chimney sample was collected on the acidic and iron-rich Capelinhos site on the Lucky Strike hydrothermal field (37°17.0’N, MAR) during the MoMARsat cruise in 2014 (http://dx.doi.org/10.17600/14000300) led by IFREMER (France) onboard R/V *Pourquoi Pas?* (Sarradin and Cannat, 2014). The sample (PL583-8) was collected by breaking off a piece of a high-temperature active black smoker using the submersible’s robotic arm and bringing it back to the surface in a decontaminated insulated box (http://video.ifremer.fr/video?id=9415). Onboard, chimney fragments were anaerobically crushed in an anaerobic chamber in an H_2_:N_2_ (2.5:97.5) atmosphere (La Calhene, France), placed in flasks under anaerobic conditions (anoxic seawater at pH 7 with 0.5 mg L^−1^ of Na_2_S and N_2_:H_2_:CO_2_ (90:5:5) gas atmosphere), and stored at 4°C.

Prior to our experiments, pieces of the hydrothermal chimney were removed from the sulfidic seawater flask, crushed with a sterile mortar and pestle in an anaerobic chamber (Coy Laboratories, Grass Lake, MI), and distributed into anaerobic tubes for use in the various experiments.

### Electrotrophic enrichment on nitrate, sulfate, and oxygen

MESs were filled with 1.5 L of an amended sterile mineral medium as previously described (Pillot *et al.*, 2019) without yeast extract and set at 80°C and pH 6.0 through on-platform monitoring. The electrode (cathode), composed of 20 cm^2^ of carbon cloth, was poised at the lowest potential before initiation of abiotic current consumption (Fig. S4) using SP-240 potentiostats and EC-Lab software (BioLogic, France). Potentials were considered for pH 6.0 and 80°C vs SHE. A potential of −590 mV vs SHE was used in the nitrate and sulfate experiments and −300 mV vs SHE in the oxygen experiment. A similar experiment at −300 mV vs SHE has been initiated in presence of sulfate (Fig. 3) to confirm the growth of electrolithoautotroph communities at lower potential, avoiding potential H_2_ production. The electrode poised as cathode served as the sole electron donor for electrotroph growth. For nitrate and sulfate experiments, the MES was supplemented with 4 mM of sodium nitrate or 10 mM of sodium sulfate, respectively. The cathodic chambers were sparged with N_2_:CO_2_ (90:10, 100 mL/min). For the oxygen experiment, the MES was sparged with N_2_:CO_2_:O_2_ (80:10:10, 100 mL/min) with initially 10% oxygen as electron acceptor. All experiments were inoculated with 8g of the crushed chimney (~0.5% (w/v)). Current consumption was monitored via the chronoamperometry method. An abiotic control without inoculation showed no current consumption during the same experiment period. CycloVoltammograms (scan rate: 20 mV/s) were analyzed using QSoas software (version 2.1). Coulombic efficiencies were calculated using the following equation:

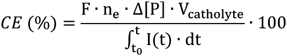

I(t): current consumed between t_0_ and t (A)
F: Faraday constant
n_e_: number of moles of electrons presents per mole of product (mol)
Δ[P]: variation of the concentration of organic product between t_0_ and t (mol.L^−1^)
V_catholyte_: volume of catholyte (L)

## Identification and quantification of organic compound production

To identify and quantify the production of organic compounds from the biofilm, samples of liquid media were collected at the beginning and at the end of the experiment and analyzed by ^1^H NMR spectroscopy. For this, 400 μL of each culture medium were added to 200 μL of PBS solution prepared in D_2_O (NaCl, 140 mM; KCl, 2.7 mM; KH_2_PO_4_, 1.5 mM; Na_2_HPO_4_, 8.1 mM; pH 7.4) supplemented with 0.5 mmol L^−1^ of trimethylsilylpropionic acid-*d*4 (TSP) as NMR reference. All the 1D ¾ NMR experiments were carried out at 300 K on a Bruker Avance spectrometer (Bruker, BioSpin Corporation, France) operating at 600 MHz for the ^1^H frequency and equipped with a 5-mm BBFO probe.

Spectra were recorded using the 1D nuclear Overhauser effect spectroscopy pulse sequence (Trd-90°-T_1_-90°-tm-90°-Taq) with a relaxation delay (Trd) of 12.5 s, a mixing time (tm) of 100 ms, and a T_1_ of 4 μs. The sequence enables optimal suppression of the water signal that dominates the spectrum. We collected 128 free induction decays (FID) of 65,536 datapoints using a spectral width of 12 kHz and an acquisition time of 2.72 s. For all spectra, FIDs were multiplied by an exponential weighting function corresponding to a line broadening of 0.3 Hz and zero-filled before Fourier transformation. NMR spectra were manually phased using Topspin 3.5 software (Bruker Biospin Corporation, France) and automatically baseline-corrected and referenced to the TSP signal (δ = −0.015 ppm) using Chenomx NMR suite v7.5 software (Chenomx Inc., Canada). A 0.3 Hz line-broadening apodization was applied prior to spectral analysis, and ^1^H-^1^H TOCSY (Bax and Davis, 1985) and ^1^H-^13^C HSQC (Schleucher et al., 1994) experiments were recorded on selected samples to identify the detected metabolites. Quantification of identified metabolites was done using Chenomx NMR suite v7.5 software (Chenomx Inc., Canada) using the TSP signal as the internal standard.

## Biodiversity analysis

Taxonomic affiliation was carried out according to Zhang et al. (2016). DNA was extracted from 1g of the crushed chimney and, at the end of each culture period, from scrapings of half of the WE and from centrifuged pellets of 50 mL of spent media. The DNA extraction was carried out using the MoBio PowerSoil DNA isolation kit (Carlsbad, CA). The V4 region of the 16S rRNA gene was amplified using the universal primers 515F (5’-GTG CCA GCM GCC GCG GTA A-3’) and 806R (5’-GGA CTA CNN GGG TAT CTA AT-3’) (Bates et al., 2011) with Taq&Load MasterMix (Promega) in triplicates and pooled together. PCR reactions, qPCR, amplicon sequencing and taxonomic affiliation were carried out as previously described (Pillot, *et al.*, 2018). The qPCR results were expressed in number of copies of 16s rRNA gene per gram of crushed chimney, per milliliter of liquid media or per cm^2^ of surface of the electrode. To analyze alpha diversity, the OTU tables were rarefied to a sampling depth of 9410 sequences per library, and three metrics were calculated: the richness component, represented by number of OTUs observed, the Shannon index, representing total biodiversity, and the evenness index (Pielou’s index), which measures distribution of individuals within species independently of species richness. Rarefaction curves (Fig. S5) for each enrichment approached an asymptote, suggesting that the sequencing depths were sufficient to capture overall microbial diversity in the studied samples. The phylogenetic tree was obtained with MEGA software v10.0.5 with the MUSCLE clustering algorithm and the Maximum Likelihood Tree Test with a Bootstrap method (2500 replications). The heatmap was obtained using RStudio software v3. The raw sequences for all samples can be found in the European Nucleotide Archive (accession number: PRJEB35427).

## Acknowledgments

This work received financial support from the CNRS-sponsored national interdisciplinary research program (PEPS-ExoMod 2016). The authors thank Céline Rommevaux and Françoise Lesongeur for taking samples during the MOMARSAT 2014 cruise, the MIM platform (MIO, France) for providing access to their confocal microscopy facility, and the GeT-PlaGe platform (GenoToul, France) for help with DNA sequencing. The project leading to this publication received European FEDER funding under Project No. 1166-39417. The authors declare no conflicts of interest.

## Supplementary material

**Supplementary Figure S1:**
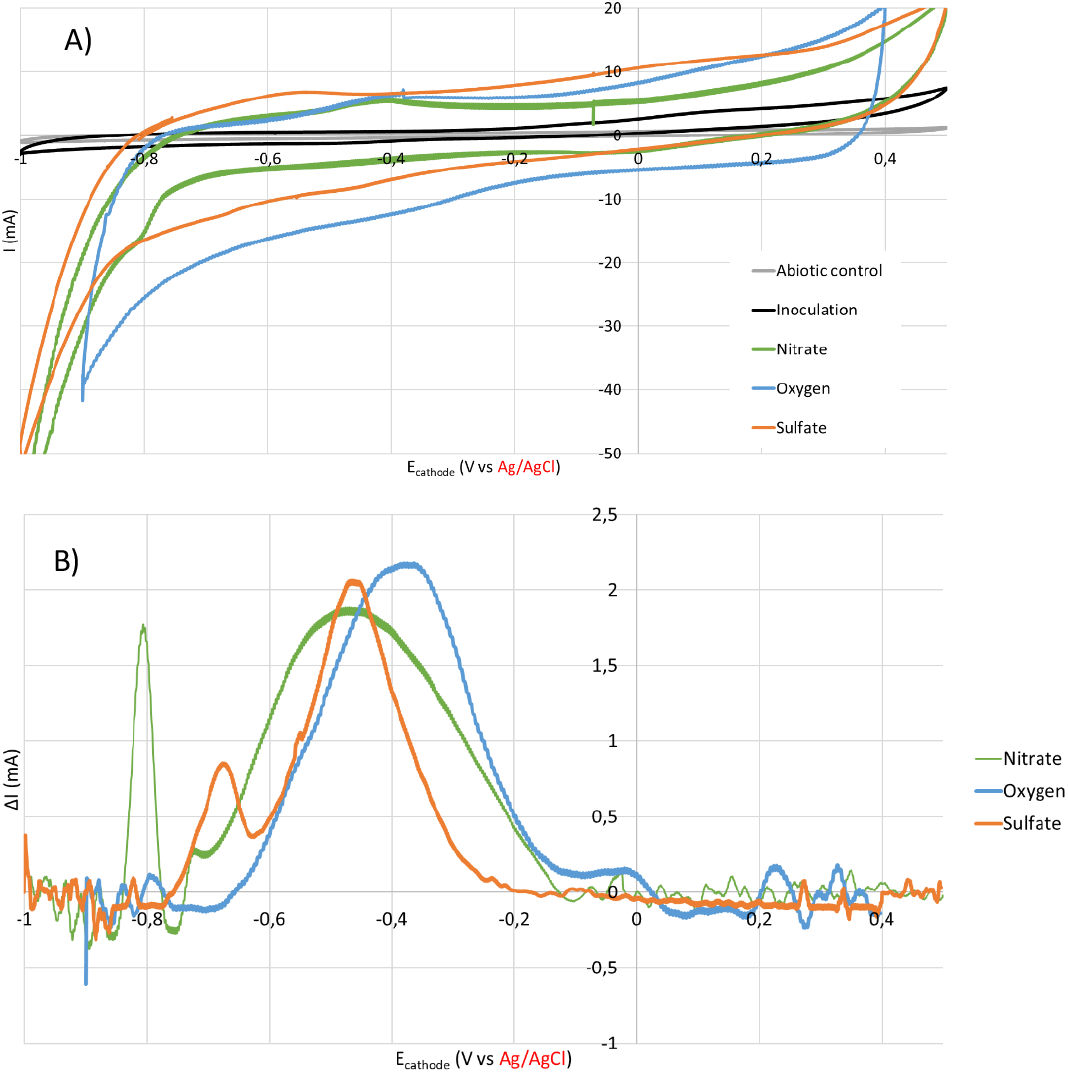
A) Cyclic Voltammograms (scan rate = 20 mV/s) of the abiotic control, and of the experiments at inoculation time and after 30 days for each condition (Nitrate, Oxygen and Sulfate). B) Reduction peaks extracted from Cyclic Voltammograms (scan rate = 20 mV/s) where the baseline have been subtracted with the software QSoas. The ΔI of reduction peaks are expressed in inversed values. Cyclovoltammetries carried out with a 3 M Ag/AgCl reference electrode (E= +0.165 V vs SHE at 80°C).

**Supplementary Figure S2.**
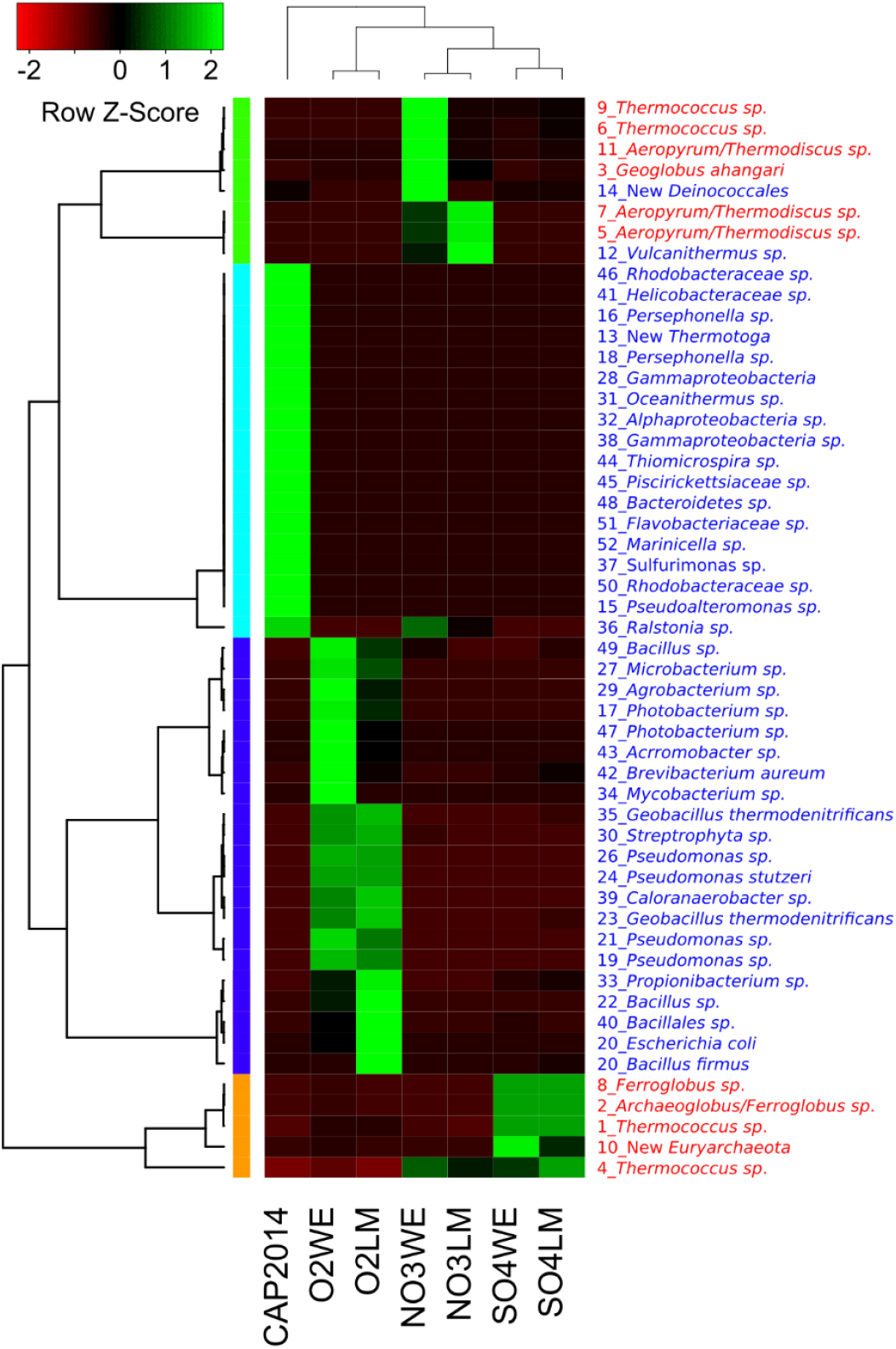
Heatmap representation of the distribution of dominant OTUs (>0.05%) over the different electron acceptors (LM: Liquid Media; WE: Working Electrode, cathode). OTUs and samples clustering were performed with centroid average method and with Pearson distance measurement method. The red taxa represent the Archaea members and blue taxa, the Bacteria.

**Supplementary Figure S3:**
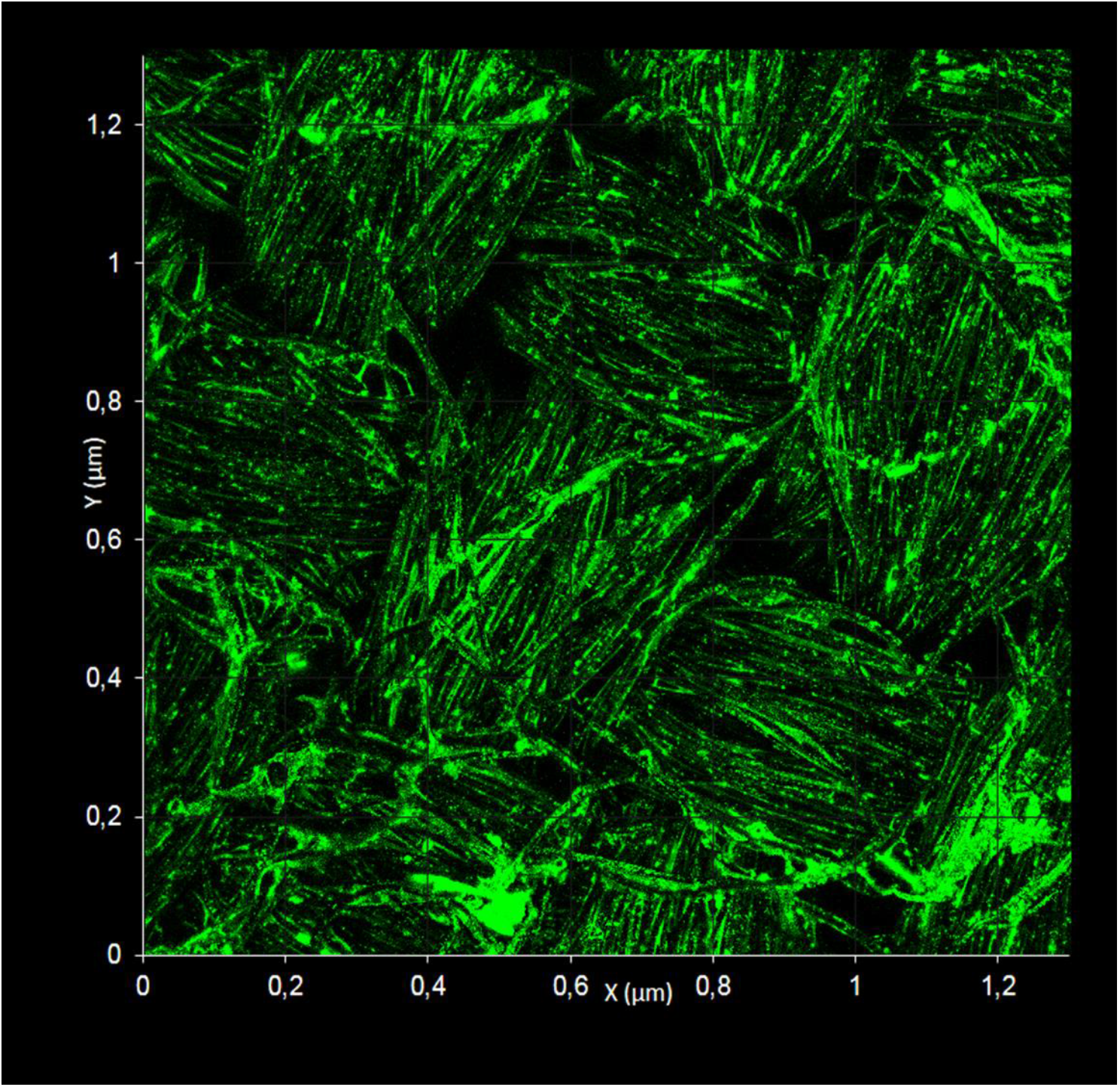
Representative confocal microscopy of the biofilm after 30 days of experiment on nitrate as electron acceptor. The green signal, corresponding to cells stained with Syto9 dye, allow to show the biofilm covering the interwoven carbon fibers.

**Supplementary Table S1:** Evolution of transient compounds (in μM) over the enrichment on Nitrate measured by NMR

| Transient compounds | Time (days) |  |  |  |  |  |  |  |  |  |  |  |  |  |  |
| --- | --- | --- | --- | --- | --- | --- | --- | --- | --- | --- | --- | --- | --- | --- | --- |
|  | 0 | 1 | 2 | 3 | 4 | 5 | 7 | 9 | 11 | 12 | 13 | 15 | 16 | 18 | 21 |
| *Benzoate like* | 5.6 | 0.0 | 29.<br>4 | 19.<br>8 | 24.<br>2 | 25.<br>5 | 54.<br>4 | 55.<br>0 | 76.<br>6 | 76.<br>2 | 94.<br>8 | 103.<br>.4 | 116.<br>.1 | 99.<br>0 | 128.<br>.4 |
| *Methanol* | 3.2 | 0.0 | 0.0 | 10.<br>8 | 16.<br>1 | 31.<br>5 | 9.0 | 36.<br>0 | 9.2 | 0.0 | 0.0 | 7.2 | 0.0 | 4.6 | 5.4 |
| *Formate* | 1.9 | 61.<br>5 | 8.4 | 54.<br>9 | 21.<br>8 | 70.<br>5 | 23.<br>4 | 63.<br>0 | 25.<br>8 | 17.<br>0 | 32.<br>6 | 23.<br>6 | 0.0 | 13.<br>4 | 22.<br>0 |
| *Cystine* | 12.<br>3 | 35.<br>5 | 30.<br>2 | 23.<br>7 | 30.<br>6 | 23.<br>6 | 17.<br>7 | 13.<br>8 | 9.6 | 7.8 | 1.2 | 2.6 | 0.0 | 0.0 | 19.<br>4 |
| *Acetoacetate* | 0.0 | 0.0 | 0.0 | 0.0 | 0.0 | 0.0 | 0.0 | 44.<br>0 | 4.6 | 15.<br>4 | 0.0 | 0.0 | 0.0 | 0.0 | 0.0 |
| *Lactate* | 6.6 | 30.<br>0 | 24.<br>0 | 15.<br>0 | 10.<br>6 | 31.<br>5 | 9.2 | 37.<br>0 | 11.<br>0 | 12.<br>2 | 17.<br>2 | 16.<br>0 | 27.<br>6 | 11.<br>0 | 13.<br>0 |
| *Threonine* | 1.9 | 11.<br>5 | 2.6 | 3.6 | 4.0 | 3.0 | 4.2 | 0.0 | 2.2 | 2.0 | 0.0 | 2.0 | 8.7 | 1.2 | 4.0 |
| *Succinate* | 1.3 | 0.0 | 2.2 | 0.0 | 0.0 | 0.0 | 0.0 | 0.0 | 0.0 | 0.0 | 0.0 | 0.0 | 0.0 | 0.0 | 1.2 |
| *Ethanol* | 2.2 | 177<br>.0 | 40.<br>6 | 37.<br>5 | 42.<br>5 | 47.<br>5 | 29.<br>8 | 86.<br>0 | 16.<br>8 | 0.0 | 33.<br>6 | 0.0 | 57.<br>9 | 0.0 | 6.4 |
| *Alanine* | 0.0 | 0.0 | 0.0 | 10.<br>8 | 12.<br>0 | 10.<br>5 | 13.<br>8 | 0.0 | 20.<br>2 | 0.0 | 54.<br>2 | 93.<br>6 | 178.<br>.5 | 179.<br>.8 | 236.<br>.6 |
| *Acetamide* | 0.0 | 0.0 | 0.0 | 0.0 | 20.<br>0 | 25.<br>5 | 48.<br>6 | 54.<br>0 | 57.<br>6 | 69.<br>2 | 39.<br>4 | 35.<br>8 | 18.<br>9 | 10.<br>6 | 6.4 |
| *2-Aminoisobutyric acid* | 4.8 | 67.<br>0 | 0.0 | 5.1 | 4.2 | 6.5 | 15.<br>2 | 28.<br>0 | 19.<br>6 | 21.<br>6 | 12.<br>0 | 13.<br>2 | 8.1 | 0.0 | 0.0 |
| *3-hydroxyisovalerate* | 4.6 | 37.<br>5 | 7.6 | 28.<br>2 | 6.6 | 34.<br>5 | 9.4 | 81.<br>0 | 13.<br>8 | 8.0 | 14.<br>4 | 11.<br>2 | 23.<br>4 | 9.8 | 14.<br>6 |

**Supplementary Figure S4:**
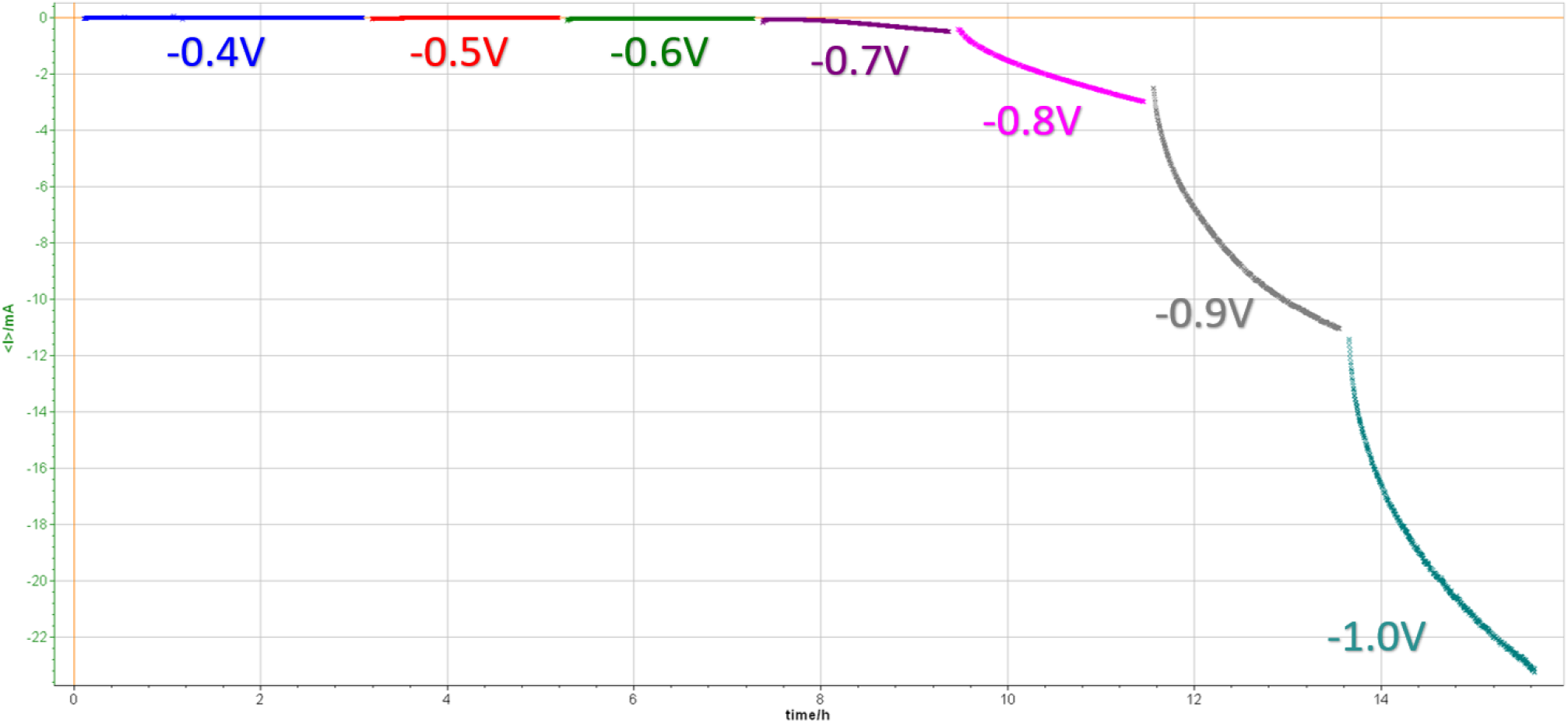
Screening of potentials in abiotic and anaerobic conditions to select the lowest potential before water electrolysis. Potential are expressed vs. SHE.

**Supplementary Figure S5.**
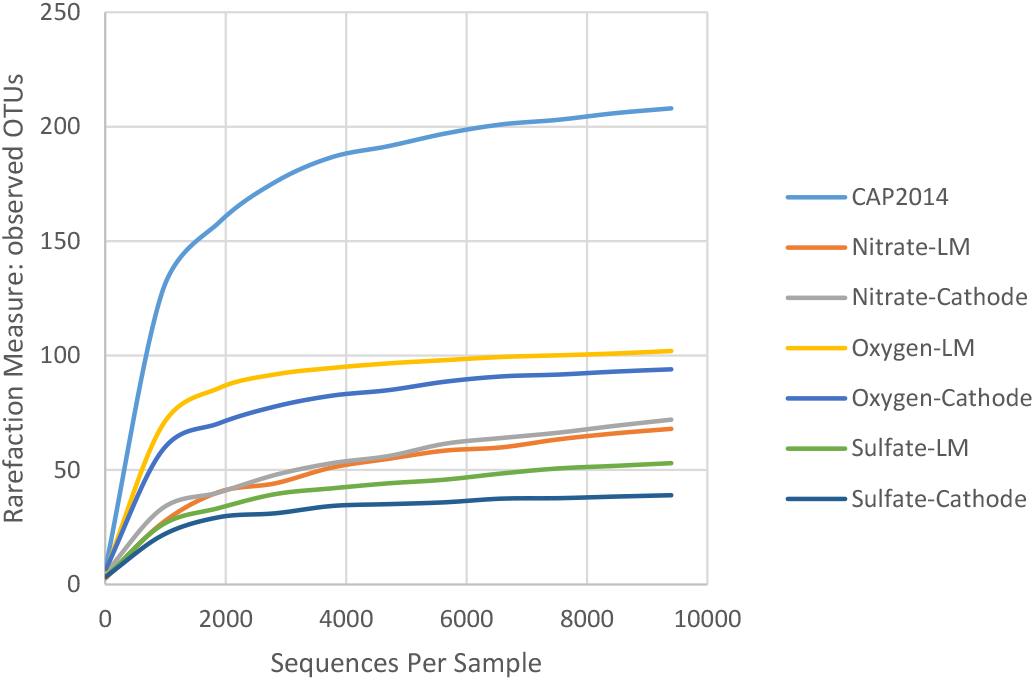
Rarefaction curves of 16S rDNA sequences for bacterial and archaeal diversities in the different samples. Curves were calculated on OTUs at 97% similarity.

**Supplementary Figure S6:**
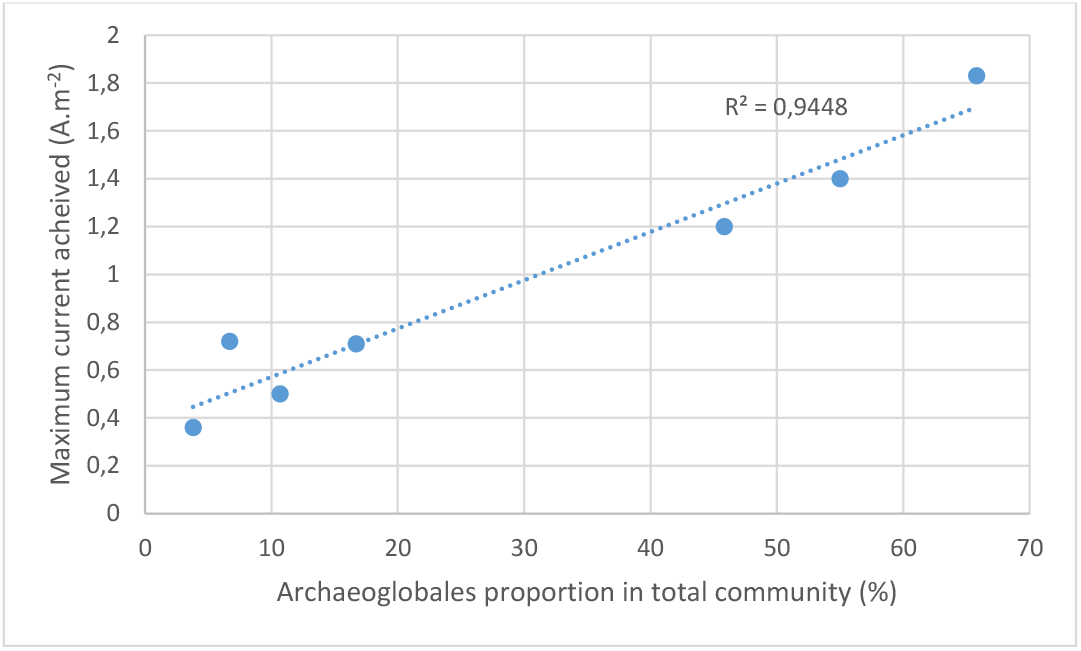
Correlation between maximum current and proportion of Archaeoglobales in the total community measured by Metabarcoding. The data were obtained from the 4 enrichments presented in this study, 2 subcultures from Nitrate enrichment and one enrichment on Fe(III)Oxide enrichment, not presented in this study.

